# Comparison of direct cDNA and PCR-cDNA Nanopore sequencing of *Escherichia coli* isolates

**DOI:** 10.1101/2024.01.23.576853

**Authors:** G Rodger, S Lipworth, L Barrett, S Oakley, DW Crook, DW Eyre, N Stoesser

## Abstract

2.

Whole-transcriptome (long-read) RNA sequencing (Oxford Nanopore Technologies, ONT) holds promise for agnostic analysis of differential gene expression (DGE) in pathogenic bacteria, including for antimicrobial resistance genes (ARGs). However, direct cDNA ONT sequencing requires large concentrations of polyadenylated mRNA, and amplification protocols may introduce technical bias. Here we evaluated the impact of direct cDNA and cDNA PCR-based ONT sequencing on transcriptomic analysis of clinical *Escherichia coli*. Four *E. coli* bloodstream infection-associated isolates (n=2 biological replicates/isolate) were sequenced using the ONT Direct cDNA Sequencing SQK-DCS109 and PCR-cDNA Barcoding SQK-PCB111.24 kits. Biological and technical replicates were distributed over 8 flow cells using 16 barcodes to minimise batch/barcoding bias. Reads were mapped to a transcript reference and transcript abundance quantified after *in silico* depletion of low abundance and rRNA genes. We found there were strong correlations between read counts using both kits and when restricting the analysis to include only ARGs. We highlighted correlations were weaker for genes with a higher GC content. Read lengths were longer for the direct cDNA kit compared to the PCR-cDNA kit whereas total yield was higher for the PCR-cDNA kit. In this small but methodologically rigorous evaluation of biological and technical replicates of isolates sequenced with the direct cDNA and PCR-cDNA ONT sequencing kits, we demonstrated that PCR-based amplification substantially improves yield with largely unbiased assessment of core gene and ARG expression. However, users of PCR-based kits should be aware of a small risk of technical bias which appears greater for genes with an unusually high (>52%)/low (<44%) GC-content.

**Impact statement:** RNA sequencing allows quantification of RNA within a biological sample providing information on the expression of genes at a particular time. This helps understand the expression of antimicrobial resistance genes (ARGs). In RNA-Seq experimental workflows extra steps of reverse transcription may be needed to generate more stable cDNA to allow for amplification by PCR if starting RNA input was low. Two current methods of long-read RNA sequencing include direct cDNA and PCR-cDNA based sequencing (Oxford Nanopore Technologies, ONT). However, few studies have compared these two methods of RNA-sequencing using clinical bacterial isolates. We therefore undertook a study to compare both kits using a methodological balanced design of biological and technical replicates of *E. coli*. Our study showed that direct cDNA and PCR-cDNA sequencing is highly reproducible between biological and technical *E. coli* replicates with very small differences in gene expression signatures generated between kits. The PCR-cDNA kit generates increased sequencing yield but a smaller proportion of mappable reads, the generation of shorter reads of lower quality and some PCR-associated bias. PCR-based amplification greatly increased sequencing yield of core genes and ARGs, however there may be a small risk of PCR-bias in genes that have a higher GC content.

**Data summary:** The transcript reads of the four sequenced *Escherichia coli* strains have been deposited in the Figshare, DOI: 10.6084/m9.figshare.25044051.

The authors confirm all supporting data (available in Figshare), code (available at: https://github.com/samlipworth/rna_methods) and protocols have been provided within the article or through supplementary data files.

## 5. Introduction

RNA sequencing (RNA-Seq) has become the leading method for transcriptome-wide analysis of differential gene expression (DGE) [1-3]. RNA-Seq can characterize all transcripts over a large dynamic range; quantify terminator efficiency and small RNAs; and measure transcriptional abundance, including operons, whilst being cost-effective [4-6]. Having a greater understanding of bacterial gene expression would provide valuable insight into the relationship between genotype and phenotype for diverse microbial functions, including antimicrobial resistance (AMR).

To date, most RNA-Seq has been based on short-read Illumina sequencing generating read lengths up to 300bp, and short-read RNA-Seq workflows and computational tools have evolved substantially [2, 5]. During library preparation for short-read sequencing however, RNA is fragmented prior to reverse transcription (RT), potentially resulting in lost information during read mapping and making distinguishing between overlapping transcripts and therefore full-length transcript analysis challenging [7, 8]. Recent advances in long-read sequencing, for example from Oxford Nanopore Technologies (ONT; hereafter referred to as “nanopore”), have removed the need for RNA fragmentation thus permitting sequencing of longer transcripts, and improved read-mapping strategies and transcript identification [7].

Nanopore RNA-Seq protocols reflect two main approaches: The first is direct sequencing of RNA molecules using the RNA-Seq kit (SQK-RNA002, upgraded to SQK-RNA004 in early 2024 for compatibility with the latest Early Access program RNA (FLO-MIN004RA) flow cell) [7-9]. The second incorporates cDNA synthesis allowing enrichment of full-length sequenced transcripts [7]. For cDNA sequencing, nanopore offer two kits – a direct cDNA sequencing kit (SQK-DCS109, evaluated in this study and hereafter referred to as the “direct” kit; replaced by SQK-LSK114 in mid-June 2023) and a kit which includes a cDNA PCR-based amplification step (SQK-PCB111.24, evaluated in this study and hereafter referred to as the “PCR” kit; pending replacement by SQK-PCB114.24 currently an early Access product, fully available in March 2024). Although direct cDNA sequencing could avoid PCR amplification bias, it also requires a high polyadenylated mRNA input (>100 ng) and the library preparation time is longer (∼5 hrs vs 1hr 45 minutes).

Bacterial RNA-Seq workflows can be challenging as messenger RNA (mRNA), typically represents <5% of RNA isolated. Most RNA extracted is ribosomal RNA (rRNA; i.e. 5S, 16S, 23S) which requires depletion to allow for sufficient, cost-effective mRNA sequencing coverage [10, 11], but poor sequence quality can result from RNA degradation during depletion. Moreover, bacterial mRNA is not naturally polyadenylated at the 3’ end, which is a pre-requisite for nanopore protocols where polyadenylated mRNA is annealed to an oligo(dT) primer for PCR-cDNA sequencing or ligated to a double-stranded oligo(dT) splint adapter in direct RNA sequencing. Given large concentrations of polyadenylated mRNA are required for nanopore sequencing, optimisation of extraction and template preparation workflows would be ideal [7, 12]. Additionally, computational tools are generally designed for eukaryotic transcriptomics or for short-read prokaryotic transcriptomics, and a bioinformatics pipeline for analysing long-read bacterial transcriptome data would be beneficial [13, 14]. Furthermore, it is essential to capture the impact of biological and experimental variability on nanopore RNA-Seq outputs, using both biological and technical replicates [15], as batch effects could result in the misidentification of differentially expressed genes [3, 13, 14, 16].

Here we compared nanopore cDNA and PCR-cDNA sequencing with a focus on the impact of AMR gene expression. We evaluated four clinical strains of *Escherichia coli*, assessing biological and technical batch effects and evidence of PCR bias with PCR-amplified cDNA libraries. Our work builds on previous experimentation on a single lab strain of *E. coli* [7].

## 6. Methods

### Isolates for testing and bacterial growth curves

We investigated four *E. coli* bloodstream infection-associated strains stored as stocks at -80°C in 10% glycerol nutrient broth; these strains had all demonstrated amoxicillin-clavulanate (co-amoxiclav) and ceftriaxone resistance using the BD Phoenix and EUCAST clinical breakpoints [17]. To evaluate bacterial growth dynamics in the absence and presence of antibiotic pressure and identify the time to mid-log phase of growth, glycerol stocks were grown on Columbia Blood Agar (CBA) overnight at 37°C. Single colonies were inoculated into 10mL of Lysogeny Broth (LB), or LB containing the breakpoint concentrations of co-amoxiclav (8 mg/L) or ceftriaxone (2 mg/L; Fig. S1), and grown at 37°C with shaking at 160rpm. The liquid suspensions were diluted and normalised to 0.05 OD_600_nm with a final volume of 200μL. The growth parameters of each isolate were measured in triplicate over 24 h at 37°C using a Tecan Spark microplate reader (Tecan Group Ltd, LifeSciences, Switzerland).

### RNA extraction

Prior to implementing the final optimised extraction and sequencing workflow described below, we analysed the outputs of several RNA extraction kits, mRNA enrichment/rRNA depletion kits, polyadenylation approaches and ONT sequencing kits, including: PureLink RNA Mini kit (Thermo Fisher Scientific, UK), MICROBExpress™ Bacterial mRNA Enrichment Kit, Poly(A) Polymerase Tailing Kit (Lucigen, UK), Direct RNA Sequencing Kit SQK-RNA002, Direct cDNA Sequencing Kit SQK-DCS109 single-plex and multiplexed using EXP-NBD104. These approaches were not used in the final workflow because of a combination of poor RNA yield post extraction, lengthy incubation times during mRNA enrichment and polyadenylation, inability to multiplex sequencing reactions (RNA002 kit), and poor overall sequencing outputs with these combinations. Details of the methodology and sequencing outputs are included in the Supplementary materials (Table S1); we have included this information so that other research teams are aware of strategies we evaluated that worked less well.

Single colonies of each of the *E. coli* strains were inoculated into 10 ml of LB and grown overnight at 37°C with shaking at 160 rpm. A 1:100 dilution of the overnight inoculum was sub-cultured in 10mL LB and grown to mid-log phase (0.5 at OD_600_nm) for RNA extraction. RNA was extracted from biological replicates (n=2 for each *E. coli* strain) on the automated KingFisher Flex platform using the MagMax™ Viral/Pathogen II Nucleic Acid Isolation Kit (Thermo Fisher Scientific, UK). Post extraction DNase treatment was performed using the TURBO DNA-*free*™ Kit (Invitrogen, UK) according to manufacturer’s instructions. Total RNA quality and integrity were assessed using the TapeStation 4200 (Agilent Technologies, USA) (Table S2) and quantified using the Broad Range RNA Qubit kit (Thermo Fisher Scientific, UK). rRNA depletion was performed using the QIAseq FastSelect 5S/16S/23S kit (Qiagen, UK) in accordance with manufacturer’s instructions with the addition of RNase inhibitor (New England BioLabs, UK) incorporated into each reaction. The addition of poly(A) tails to mRNA transcripts was performed using the Poly(A) Polymerase (New England BioLabs, UK) and included the addition of RNase inhibitor (New England BioLabs, UK) in each reaction. A final clean-up of mRNA using RNAClean XP beads (Beckman Coulter) was performed prior to library preparation. mRNA was checked pre and post poly(A)tail addition using an Agilent RNA 6000 Nano kit run on a 2100 BioAnalyser (Agilent Technologies, USA) (Fig. S2). Quantifications were performed using the High Sensitivity RNA Qubit kit (Thermo Fisher Scientific, UK). To alleviate any possible degradation of RNA repeated free/thawing cycles were avoided by completing the steps from extraction to cDNA synthesis in a single day.

### RNA Library preparation and sequencing

Two sequencing kits were selected for comparison in the final workflow: Direct cDNA Sequencing SQK-DCS109 and PCR-cDNA Barcoding SQK-PCB111.24 with libraries prepared in accordance with manufacturer’s instructions. Our experimental design tested differential expression between *E. coli* (n=4 strains, n=2 biological replicates/strain) using a balanced block design [15]. Technical replicates (n=2 per biological replicate) were multiplexed (n=16 replicates/flow cell in total) across 4 flow cells per sequencing kit (i.e. n=8 flow cells in total); Fig.1). Nanopore libraries were sequenced using R9.4.1 (FLO-MIN106) flow cells on a GridION with MinKNOW software version 21.11.7 and basecalled using Guppy (version 6.1.5) for the maximum 72 hour run time.

**Figure 1.**
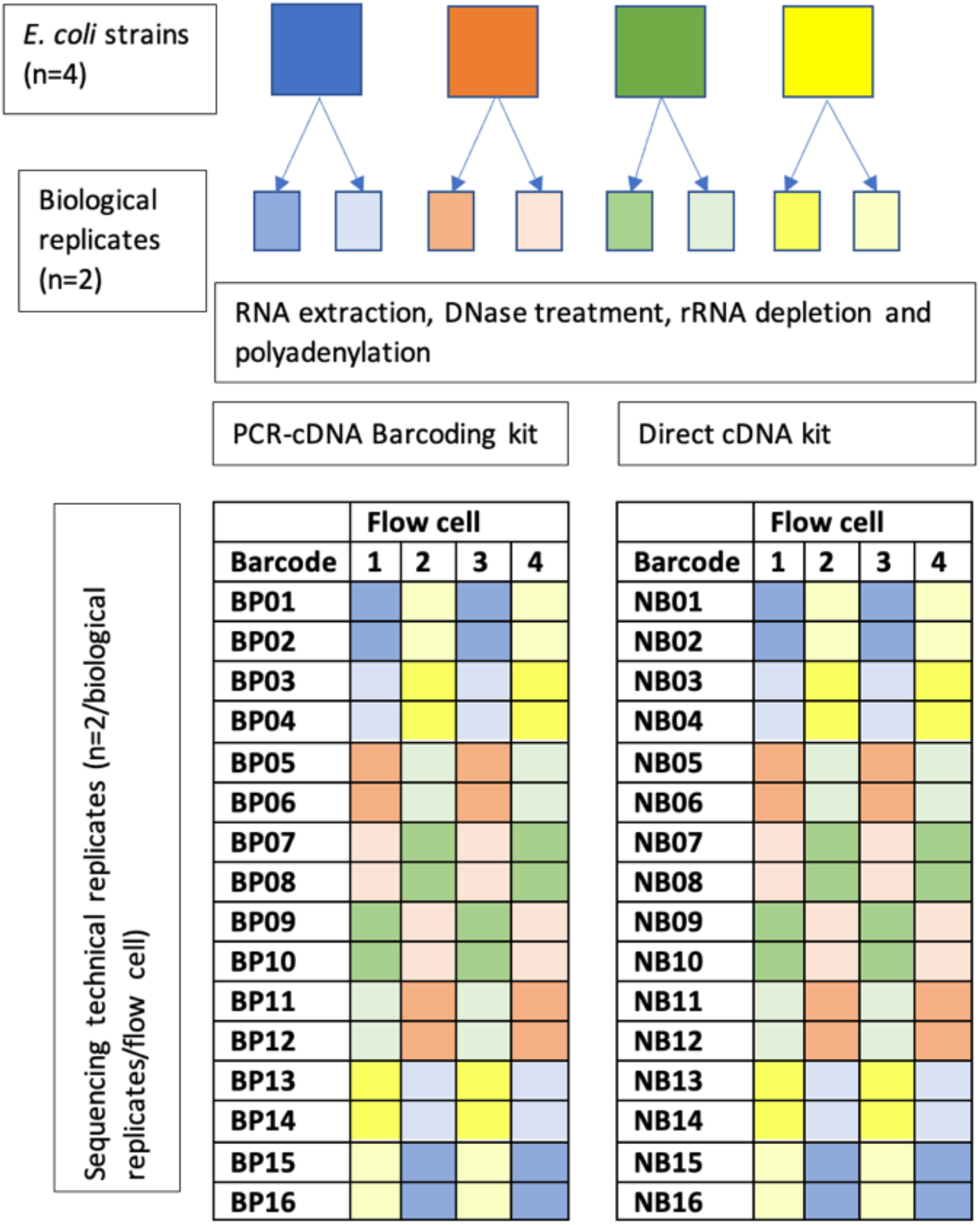
experimental design illustrating number of biological and technical replicates sequenced per flow cell and per kit.

### DNA extraction and sequencing

Short-read sequencing data (Illumina HiSeq) was created as part of a previous project (NCBI BioProject accession: PRJNA604975). DNA was extracted for nanopore sequencing using the automated KingFisher Flex platform using the MagMax™ Viral/Pathogen II Nucleic Acid Isolation Kit (Thermo Fisher Scientific, UK). DNA was sequenced on an ONT GridION using the rapid barcoding kit (SQK-RBK004) R9.4.1 (FLO-MIN106) flow cells with MinKNOW software version 21.11.7 for 48 hours run time and basecalled using Guppy (version 5.1.13).

### Sequence analysis

Raw fast5 files were basecalled and demultiplexed using Guppy (version 6.1.5). Sequence length and quality statistics were extracted using NanoStat (v1.4.0). Hybrid assemblies were created using Unicycler (--mode bold) and subsequently annotated with Bakta (v1.6.1) to create transcript references. Reads were then mapped to the transcript reference using Minimap2 (-ax splice, version 2.17-r941) [18]. In addition to the efforts to deplete rRNA in the laboratory workflow described above, reads mapping to rRNA coding sequences were discarded from further analysis. Transcript abundance was quantified using Salmon (version 1.8.0) [19]. AMR genes were identified using AMRFinder (version 3.11.2) [20]. NanoPlot was used to assess the relationship between read quality and length after randomly downsampling sequenced data to ∼250Mb using Rasusa for computational feasibility.

#### Statistical analysis

EdgeR (v3.42.2) was used to fit quasi-likelihood negative binomial generalised log-linear models (glmQLFit) to this count data (after filtering using filterByExpr min.count=1) to compare e.g. expression between direct and PCR kits. Distributions (of e.g. read quality scores) between groups were compared using Kruskal-Wallis tests and correlations using Spearman’s coefficient. Proportions between groups in 2×2 tables were compared using Fisher’s exact tests. All statistical analysis was performed in R version 4.2.1 [21].

## 7. Results

### Description of isolates

The four isolates used in this study were identified as belonging to STs 131 (A,C), 1193 (D) and an unclassified ST (B). A total of 36 AMR genes (ARGs) in total were identified: 12 in isolate A, 7 in isolate B, 7 in isolate C and 10 in isolate D (Table 1). The reference transcript sizes for each isolate were: Isolate A - 5,425 coding sequences (4,758,606bp), Isolate B - 5,570 coding sequences (4,830,938bp), Isolate C - 5,523 coding sequences (4,809,970bp), and Isolate D - 5,191 coding sequences (4,564,879bp).

**Table 1.** Summary of isolate assemblies and ARGs detected.

| Isolate | Contig | ARGs |
| --- | --- | --- |
| A | Chromosome - 5,216,957bp | 2 copies *bla*<sub>CTX-M-15</sub><br>1 copy *aac(3)-Ile*, *catB3*, *bla*<sub>OXA-1</sub>, *aac(6')-Ib-cr5*, *tet(A)*, *bla*<sub>TEM-1</sub>, *dfrA14*, *sul2*, *aph(3'')-Ib*, *aph(6)-Id* |
|  | Plasmid 1 - 113,482bp | - |
|  | Plasmid 2 - 4,236bp | - |
| B | Chromosome - 5,174,320bp | - |
|  | Plasmid 1 - 138,864bp | *tet(B)*, *sul1*, *aadA5*, *dfrA17* |
|  | Plasmid 2 - 97,597bp | *tet(B)*, *aph(3'')-Ib*, *aph(6)-Id* |
|  | Plasmid 3 - 7940bp | - |
|  | Plasmid 4 - 5,631bp | - |
|  | Plasmid 5 - 4,237bp | - |
|  | Plasmid 6 - 4,072bp | - |
|  | Plasmid 7 - 2,101bp | - |
|  | Plasmid 8 - 1,552bp | - |
| C | Chromosome - 5,113,180bp | - |
|  | Plasmid 1 - 116,873bp | *dfrA17*, *tet(A)*, *aph(6)-Id*, *aph(3'')-Ib*, *sul2* |
|  | Plasmid 2 - 87,246bp | - |
|  | Plasmid 3 - 71,559bp | - |
|  | Plasmid 4 - 8,957bp | *mph(A)*, *bla*<sub>CTX-M-27</sub> |
|  | Plasmid 5 - 5,631bp | - |
|  | Plasmid 6 - 5,221bp | - |
|  | Plasmid 7 - 5,166bp | - |
|  | Plasmid 8 - 4,082bp | - |
|  | Plasmid 9 - 1,718bp | - |
| D | Chromosome - 4,959,734bp | *bla*<sub>CTX-M-15</sub>, *aac(6')-Ib-cr5*, *bla*<sub>OXA-1</sub>, *catB3*,<br>*aac(3)-Ile* |
|  | Plasmid 1 - 92,583bp | *sul2*, *aph(3'')-Ib*, *aph(6)-Id*, *sul1*, *dfrA5* |
|  | Plasmid 2 - 79,495bp | - |

### The PCR kit produces a greater sequencing yield but with shorter read lengths and lower quality scores

The total data yield averaged across 4 flow cells after 72 hours and multiplexing 16 barcoded samples was 1.8Gb and 11.0Gb for the direct and PCR kits, respectively. However, median read lengths produced by the direct kit were longer than those produced by the PCR kit (501 bp [IQR: 390-603] versus 318 bp [IQR: 293-400]; p<0.001) (Fig. S3). Read quality (Fig. S4) was broadly comparable between kits, though slightly higher Q-score values were obtained for the direct versus PCR kit (median Q-score: 12.0 [IQR: 10.3-13.9] vs 11.2 [IQR: 10.4-12.1], p<0.001).

### Mappable reads are longer and higher quality and represent a greater proportion of total reads in the direct kit

Overall, the percentage of reads that could be mapped to the respective reference transcript was higher for the direct vs the PCR kit (median mapped percentage: 47% [IQR: 41-55%] vs 85% [IQR: 77-88%], p<0.001). Using the direct kit, there was no difference in the % of reads that could be mapped between isolates (Isolate A median: 83% [IQR: 82-88] reads mapped, Isolate B median: 86% [IQR: 80-87] reads mapped, Isolate C median: 83% [IQR 76-89] reads mapped, Isolate D median: 85% [IQR: 82-88] reads mapped; p=1.00), however this was not the case for the PCR kit (Isolate A median: 48% [IQR: 42-55] reads mapped, Isolate B median: 46% [IQR: 39-51] reads mapped, Isolate C median: 61 [IQR: 48-64] reads mapped, Isolate D median: 41 [IQR 38-47] reads mapped; p=0.02, Fig. S5).

For both the direct and PCR kits, reads that could be mapped were longer (direct mapped median read length: 510 bases [IQR: 388-608] versus direct unmapped median read length: 361 bases [IQR: 298-466], p<0.001 and PCR mapped median read length: 437 bases [IQR: 342-569] versus unmapped median read length: 301 bases [IQR: 283-322], p<0.001). They were also of higher quality (direct mapped median Q score: 12.4 [IQR: 10.4-14.3] versus direct unmapped median Q score: 11.2 [IQR: 10.1-12.8], p<0.001 and PCR mapped median Q score: 11.8 [IQR 10.9-12.7] versus unmapped median Q score: 11.0 [IQR: 10.2-11.8], p<0.001) than those that could not be mapped (Fig. S6).

### Read counts are strongly correlated for biological replicates and kits

For n=3786/3213/3381/3998 (in Isolates A-D respectively) individual coding sequences evaluated, read counts were highly correlated between biological replicates for all isolates for both the direct and PCR kits (R^2^ range: 0.90-0.98, p<0.001, Fig. 2). The correlations between technologies were similarly strongly positive (R^2^ range: 0.93-0.96, p<0.001) although we observed that these correlations were weaker for coding sequences with a high GC content (here defined as GC content >52%) or low GC content (defined as GC content <44%) (Fig. 3). Strong correlations between read counts were also seen amongst flow cells for biological replicates sequenced using the same kit (Fig. S7). Restricting only to ARGs also revealed very strong correlations between read counts for biological replicates and between kits (R^2^ range: 0.93-0.99, p<0.001).

**Figure 2.**
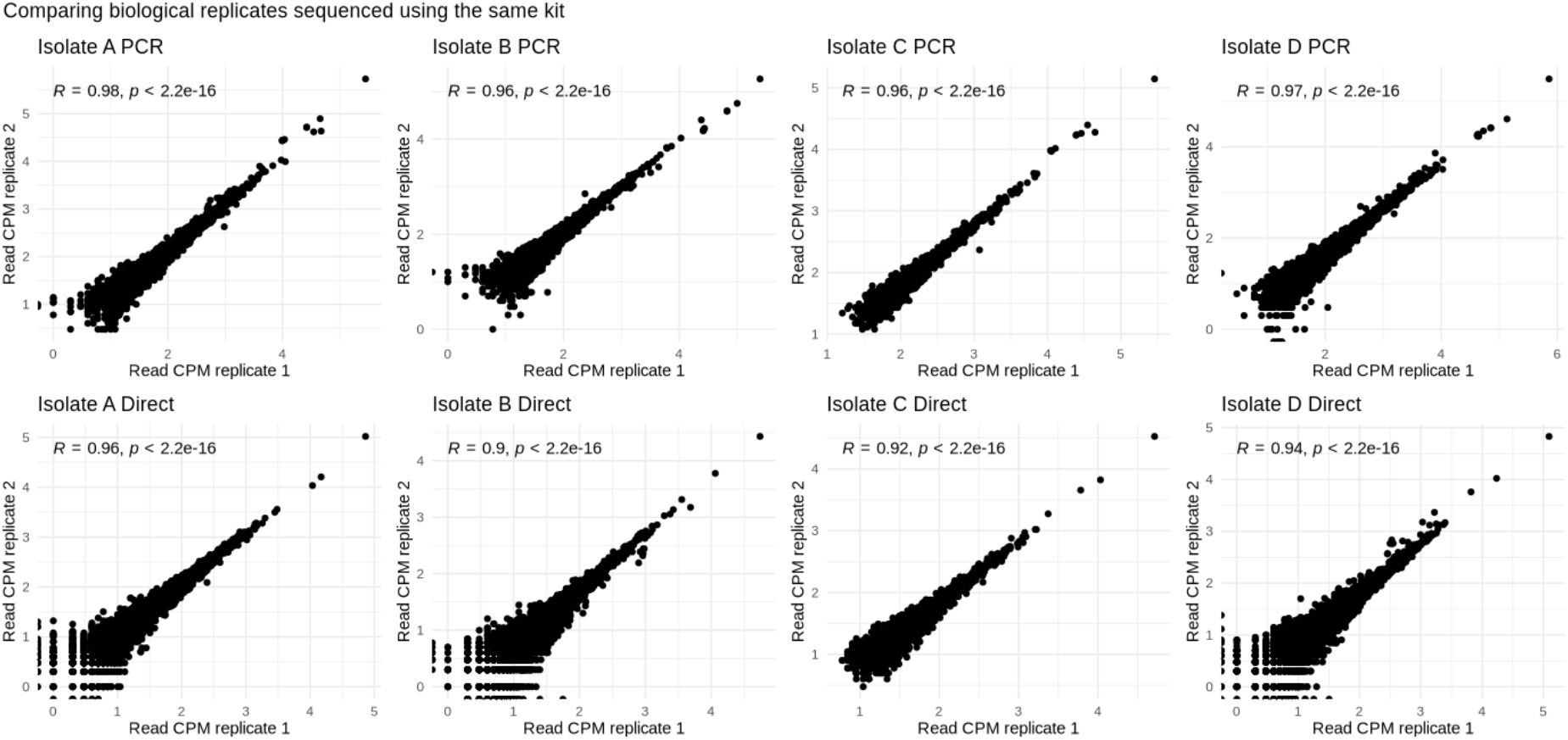
Read count correlations when comparing: (upper row) biological replicates sequenced with the PCR kit; (lower row) biological replicates sequenced using the direct kit. CPM - counts per million. Spearman correlation coefficients and p-values are shown within each plot.

**Figure 3.**
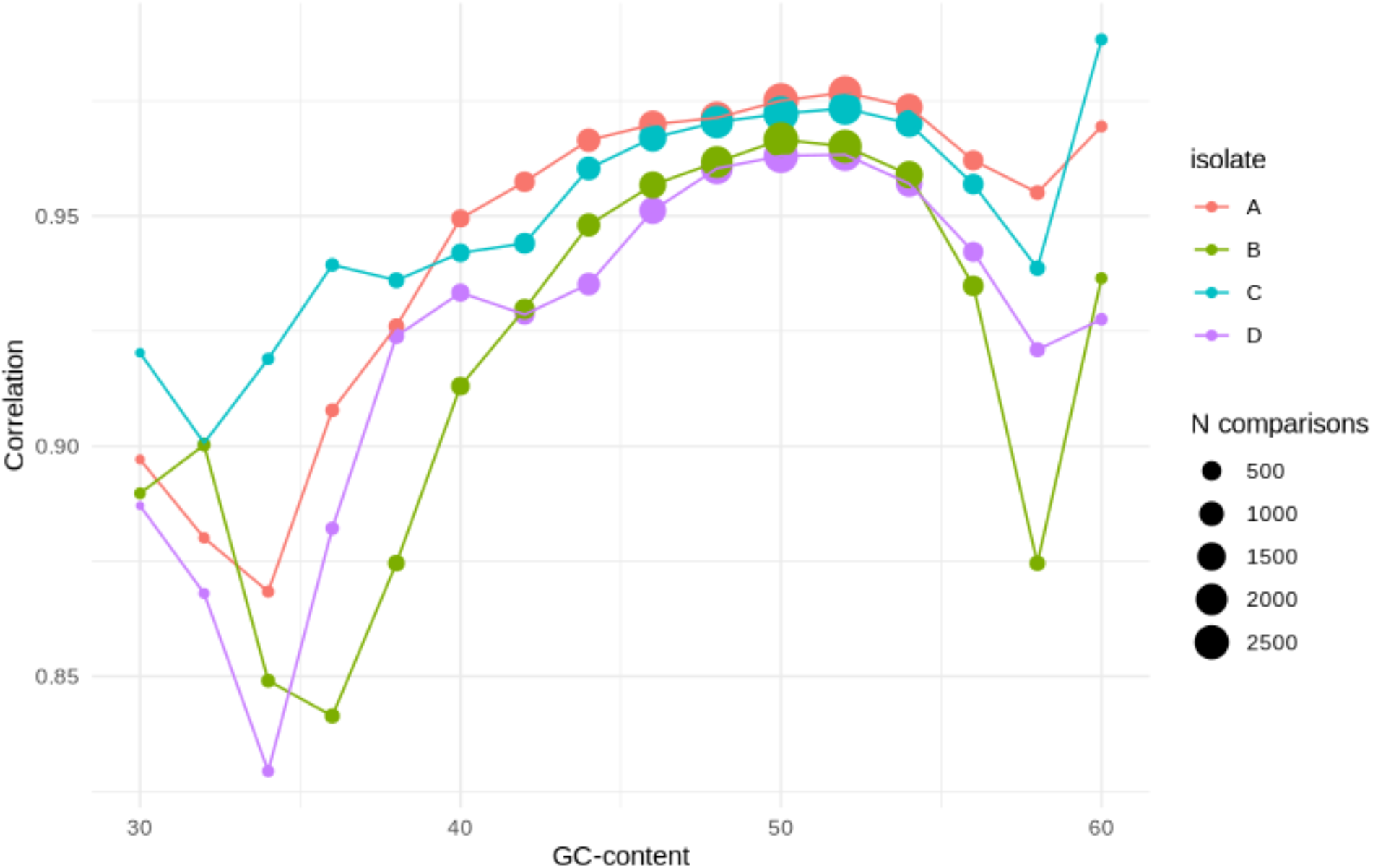
Read count correlation between the PCR and direct kits for genes with different %GC-content.

### Differences in gene “expression” by sequencing method

Across the four isolates, of those annotated coding sequences with reads mapping to them which were not rRNA and after correction for multiple comparisons, 678/14,378 (4.7%) coding sequences were observed to have significantly different “expression” between the direct and PCR kits (Fig. 4). In comparison only 31/14,378 (0.2%) coding sequences were significantly differentially expressed between biological replicates of the same isolate using the same kit. In the PCR based kit, transcripts mapping to “over-expressed” coding sequences were significantly shorter (359bp (IQR 320-576) vs non over-expressed 438bp (344-569), p<0.001). There was no difference in the proportion of plasmid (13/317, 4.1%) and chromosomal genes (539/14,122, 3.8%) that were significantly differentially expressed between the direct and PCR kits (p=0.91). Similarly, none of the 36 ARGs in the analysis were significantly differentially expressed between kits.

**Figure 4.**
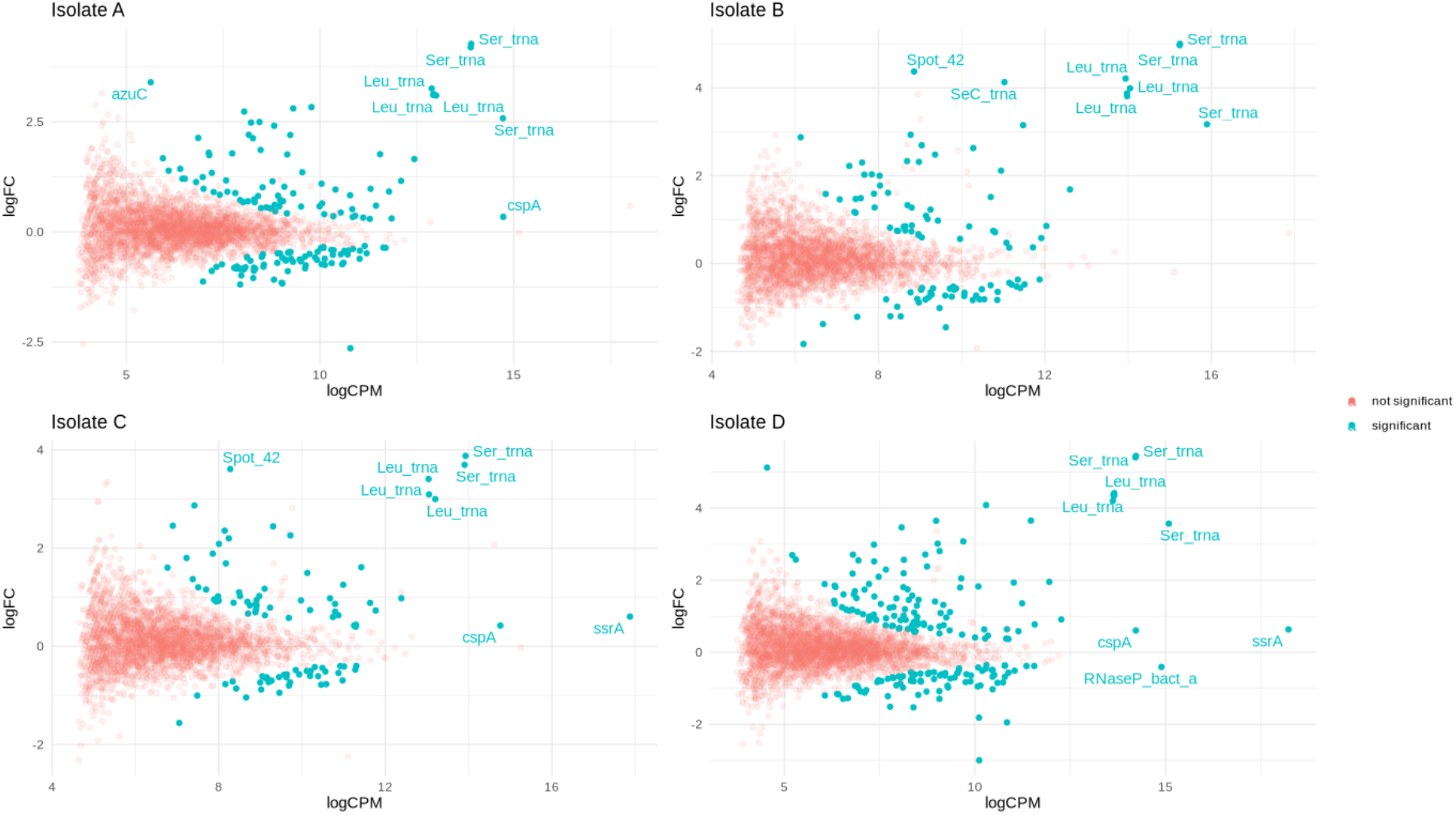
Gene “expression” differences observed between sequencing methods. For each isolate (A/B/C/D), the log count per million (logCPM) for each gene is plotted against the log-fold change (logFC) between PCR and direct kits. Genes with significant differences between kits after adjustment for multiple comparisons are shown in blue. Genes are annotated if they are significantly differentially expressed between the kits and are in the top 5 ranked genes for either logCPM or logFC.

## 8. Discussion

The ability to sequence long mRNA transcripts using nanopore sequencing offers the potential to identify and quantify transcripts in a single assay which could help to evaluate the relationship between genotype and phenotype for key clinically relevant traits (e.g. AMR). In this study, using clinical *E. coli* isolates carrying relevant AMR genes, we describe a laboratory and bioinformatic workflow for nanopore RNA-seq and show that gene expression counts are highly correlated between biological replicates and flow cells, although there is some evidence of significant differences in expression signatures generated for ∼5% of genes depending on the kit. Notably many coding sequences that were differentially “expressed” for experiments using the PCR vs direct kits were tRNAs (Fig.4), consistent with a known preference for PCR to amplify shorter fragments in a cDNA library mix of variable fragment lengths [1, 7]. However, expression signatures in our dataset were similar for both kits, suggesting that PCR-based nanopore RNA-Seq would be a robust way of defining this for specific research questions, bearing in mind this potential methodological bias.

Our findings are similar to a previous study evaluating direct and PCR-based nanopore RNA-Seq that used a single relatively antibiotic susceptible reference isolate [7], and also demonstrated highly concordant results between methods except for genes with unusually high or low GC-content (i.e. <46/>54%). Here, we extend these findings and show that the PCR kit remains suitable for use with multi-drug-resistant strains containing multiple plasmid-based ARGs. To our knowledge this study provides the first such data supporting the idea that this methodology is suitable for quantitative transcription analysis in these priority clinical isolates.

The inclusion of only four isolates, whilst to our knowledge representing the largest evaluation of the use of nanopore RNA-seq for clinically relevant *E. coli* strains, is nevertheless a significant limitation. In common with existing literature in this area and primarily due to the very high costs associated with this technology, our study lacks negative controls and so we cannot determine the degree to which the “kitome” may explain some of the variation observed. Another limitation is that our analysis only captures RNA expression in a single environmental state (growth in antibiotic-free culture medium) which likely does not represent the eventual use case (quantification of gene expression in the presence of antibiotics). The PCR step in the PCR-based kit, when following manufacturer’s instructions, has a preference is to overamplify short fragments, especially at a higher number of cycles [1, 7]. Therefore, optimisation of this reaction is warranted to try to reduce any biases. Finally, the kits and flow cells used in this experiment have been recently superseded by newer flow cells and sequencing chemistries (RNA flow cell, flow cell version R10.4.1 with v14 sequencing chemistry) and direct RNA, direct cDNA and cDNA PCR-based amplification kits; although it seems unlikely that this would affect the findings of this study, repeat characterisation using our workflow as a framework would be warranted for these and future upgrades, which occur frequently.

In summary, we demonstrate that nanopore RNA sequencing appears to be highly reproducible between biological and technical replicates, with minimal difference in gene expression signatures generated when using the PCR versus the direct kit. Potential users need to weigh up the benefits associated with the PCR kit of greatly increased sequencing yield and therefore analytical feasibility with the potential drawbacks, namely a smaller proportion of mappable reads, the apparent generation of shorter reads of a lower quality, and a small risk of PCR-associated bias.

## 10. Author statements

### 10.1 Author contributions

GR, SL and NS designed the study. GR performed all experiments. SL mapped, annotated and analysed the transcriptome data and performed data upload. GR, SL and NS drafted the manuscript. All authors contributed to the editing and a critical analysis of the manuscript.

### 10.3 Funding information

This research was funded by a John Fell Fund award to NS and DWE (grant number: 0008776). This study was also supported by the National Institute for Health Research (NIHR) Health Protection Research Unit in Healthcare Associated Infections and Antimicrobial Resistance (NIHR200915), a partnership between the UK Health Security Agency (UKHSA) and the University of Oxford. Additionally, the study was supported by the National Institute for Health Research (NIHR) Oxford Biomedical Research Centre (BRC). The views expressed are those of the author(s) and not necessarily those of the NIHR, UKHSA, the Department of Health and Social Care or the NHS.

NS is an Oxford Martin Fellow.

### 10.4 Ethical approval

No ethical approval was required for this study.

### 10.5 Consent for publication

Not applicable.

## 10.6 Acknowledgements

We are grateful to the John Radcliffe Microbiology laboratory and team who facilitate routine collection of clinical bacterial isolates.

