## Supplementary materials for "Comparison of direct cDNA and PCR-cDNA Nanopore sequencing of *Escherichia coli* isolates"

**Supplementary methods**

**Figure S1.** Growth curve analysis of all four strains (in triplicate) in breakpoint MIC concentrations of ceftriaxone (2 mg/L), co-amoxiclav (8 mg/L) and LB broth measured over 24 hours at 37°C, with LB broth incubated as the control. These data were used to determine the time to harvest each strain in mid-log phase for RNA extraction.

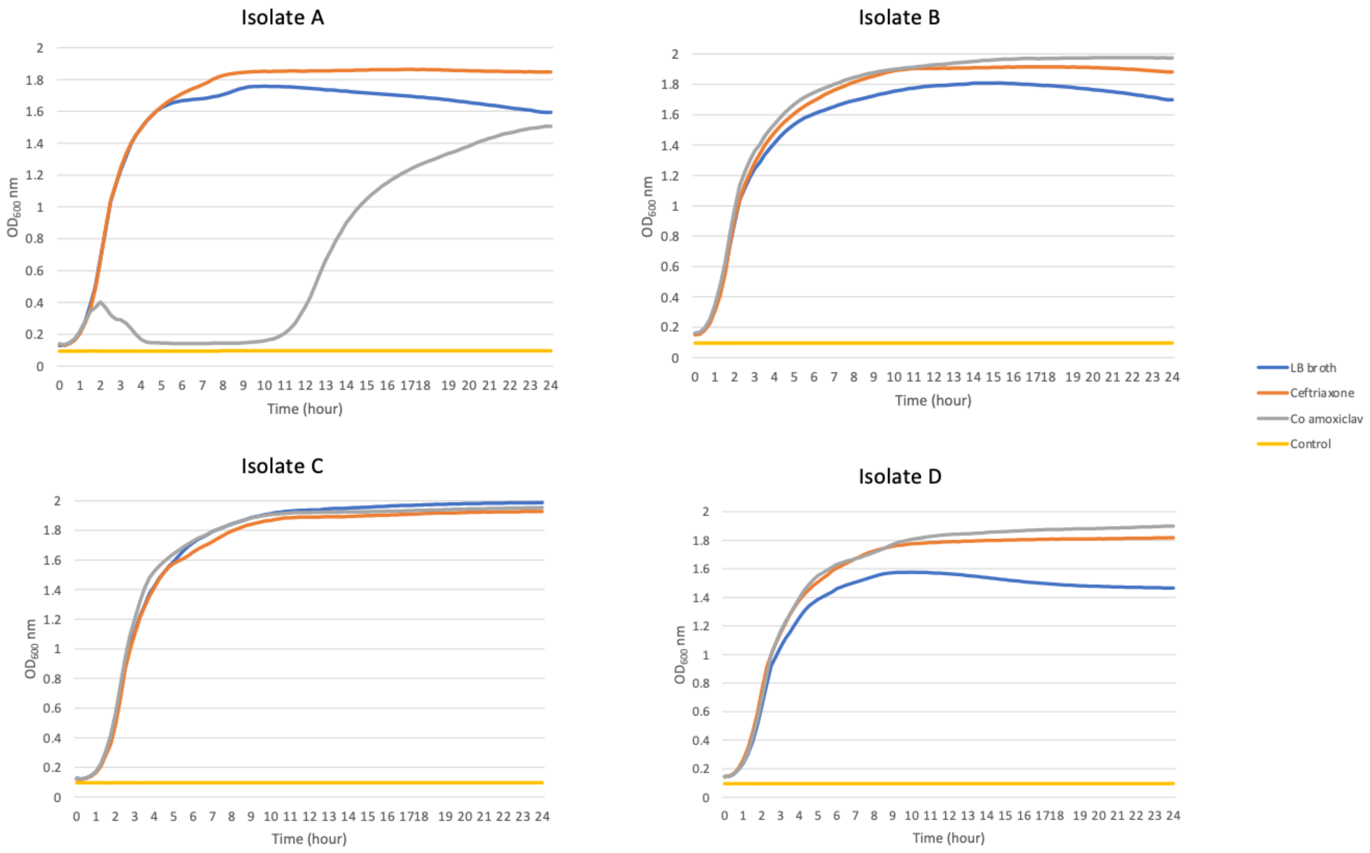

**Table S1. Different approaches trialled prior to implementation of the optimal approach and sequencing outputs.** These approaches were not used because of a combination of poor RNA yield post extraction, lengthy incubation times during mRNA enrichment and polyadenylation, unable to multiplex sequencing reactions (RNA002 kit) and overall poor sequence output.

| **RNA extraction** | **mRNA enrichment or rRNA depletion kit** | **Polyadenylation kit** | **Sequencing approach** | **ONT kit** | **Strain** | **Run number. Sample number** | **Total bases** | **rRNA reads** | **rRNA bases** | **mRNA/ rRNA ratio** | ***E. coli* mRNA reads (n)** | ***E. coli* mRNA bases (n)** | **unmapped reads** | **unmapped bases** | **Notes** | **Reference** |
|---|---|---|---|---|---|---|---|---|---|---|---|---|---|---|---|---|
| PureLink RNA Mini kit | MICROBExpress™ Bacterial mRNA Enrichment Kit | Poly(A) Polymerase Tailing Kit | Direct RNA sequencing | SQK-RNA 002 | D | 1 | 4,567 | 3 | 1,568 | 0 | 0 | 0 | 9 | 2,999 | Output mapping to non-*E. coli* (i.e. 'kit-ome') | [1] |
| | | | | | D | 2 | 7,380,461 | 1 | 913 | 0 | 0 | 0 | 9,252 | 7,379,548 | Output mapping to non-*E. coli* (i.e. 'kit-ome') | |
| | | | | | D | 3 | 65,346,291 | 43,710 | 39,277,663 | 0.021655309 | 1647 | 869,397 | 33,964 | 25,199,231 | aac-3-IIa and blaCTX-M genes detected | |
| | | | | | D | 4 | 11,114,341 | 30 | 15,483 | 0.634092735 | 76 | 26,831 | 15,483 | 11,072,027 | no resfinder genes detected | |
| | | | | | D | 5 | 3,044,399 | 4 | 897 | 0.672866521 | 3 | 1,845 | 4,089 | 3,041,657 | poor quality flowcell | |

| | | | | | | | | | | | | | | | |
|---|---|---|---|---|---|---|---|---|---|---|---|---|---|---|---|
| | | | | | D | 6 | 2,257,324 | 121 | 103,882 | 0.856876558 | 1,415 | 621,939 | 2,136 | 1,531,503 | aac-3-IIa and blaCTX-M genes detected |
| | | | | | D | 7 | 5,921,810 | 51 | 20,426 | 0.691030101 | 310 | 45,684 | 11,610 | 5,855,700 | no resfinder genes detected |
| MagMax™ Viral/Pathogen II Nucleic Acid Isolation Kit | QIAseq FastSelect 5S/16S/23S kit | *E. coli* Poly(A) Polymerase | Direct RNA sequencing | SQK-RNA002 | D | 8 | 137,353,879 | 31,491 | 13,867,854 | 0.56 | 21,918 | 7,828,855 | 277,766 | 115,657,170 | Total RNA from KF |
| | | | | | | 9 | 21,125,739 | 8,113 | 5,679,799 | 0.08 | 684 | 457,335 | 26,130 | 14,988,605 | polyadenylated |
| MagMax™ Viral/Pathogen II Nucleic Acid Isolation Kit | QIAseq FastSelect 5S/16S/23S kit | Poly(A) Polymerase Tailing Kit | Direct cDNA sequencing | SQK-DCS109 With EXP-NBD104 | D | 10.1 | 89,079,125 | 42,813 | 72,223,452 | 0.22 | 2,950 | 15,674,984 | 1,269 | 1,180,689 | sample direct from KF extraction |
| | | | | | | 10.2 | 206,653,753 | 95,611 | 192,760,506 | 0.06 | 5,496 | 12,011,067 | 2,648 | 1,882,180 | sample direct from KF extraction |
| | | | | | | 10.3 | 117,850,299 | 85,596 | 94,662,096 | 0.16 | 27,349 | 15,096,266 | 18,080 | 8,091,937 | Polyadenylated |

| | | | | | | | | | | | | | | | |
|---|---|---|---|---|---|---|---|---|---|---|---|---|---|---|---|
| | | | | | | 10.4 | 62,581,978 | 13,838 | 19,656,939 | 1.25 | 55,603 | 24,633,423 | 47,672 | 18,291,616 | rRNA depleted + polyadenylated |
| | | | | | | 10.5 | 41,676,647 | 16,601 | 24,101,650 | 0.66 | 3,296 | 15,884,314 | 2,647 | 1,690,683 | mRNA enriched + polyadenylated |
| MagMax™ Viral/Pathogen II Nucleic Acid Isolation Kit | QIAseq FastSelect 5S/16S/23S kit | *E. coli* Poly(A) Polymerase | Direct cDNA sequencing | SQK-DCS109 | D | 11 | 125,781,561 | 663 | 810,573 | 133 | 224,848 | 107,573,931 | | | |
| | | | | | | 12 | 152,628,931 | 488 | 244,024 | 533 | 302,700 | 130,052,511 | | | |
| | | | | | | 13 | 65,310,354 | 221 | 103,915 | 553 | 135,314 | 57,461,226 | | | |
| | | | | | | 14 | 430,791,822 | 2,755 | 3,856,749 | 95 | 566,642 | 366,044,621 | | | |

**Table S2.** RNA Integrity numbers (RIN) of total RNA and DNase treated RNA assessed using the TapeStation 4200 (Agilent Technologies, USA)

| Isolate (biological replicate) | RIN number (total RNA) | RIN number (DNase treated RNA) |
|---|---|---|
| A (1) | 7.4 | 7.4 |
| A (2) | 7.2 | 7.0 |
| B (1) | 7.0 | 7.1 |
| B (2) | 7.4 | 7.1 |
| C (1) | 7.4 | 7.7 |
| C (2) | 7.2 | 7.3 |
| D (1) | 7.4 | 7.3 |
| D (2) | 7.5 | 7.8 |

**Figure S2.** Electropherograms of mRNA from strains following rRNA depletion and polyadenylation (PA) with marker peaks at 25 and 4000 nt measured using the Agilent RNA 6000 Nano kit run on a 2100 BioAnalyser (Agilent Technologies, USA)

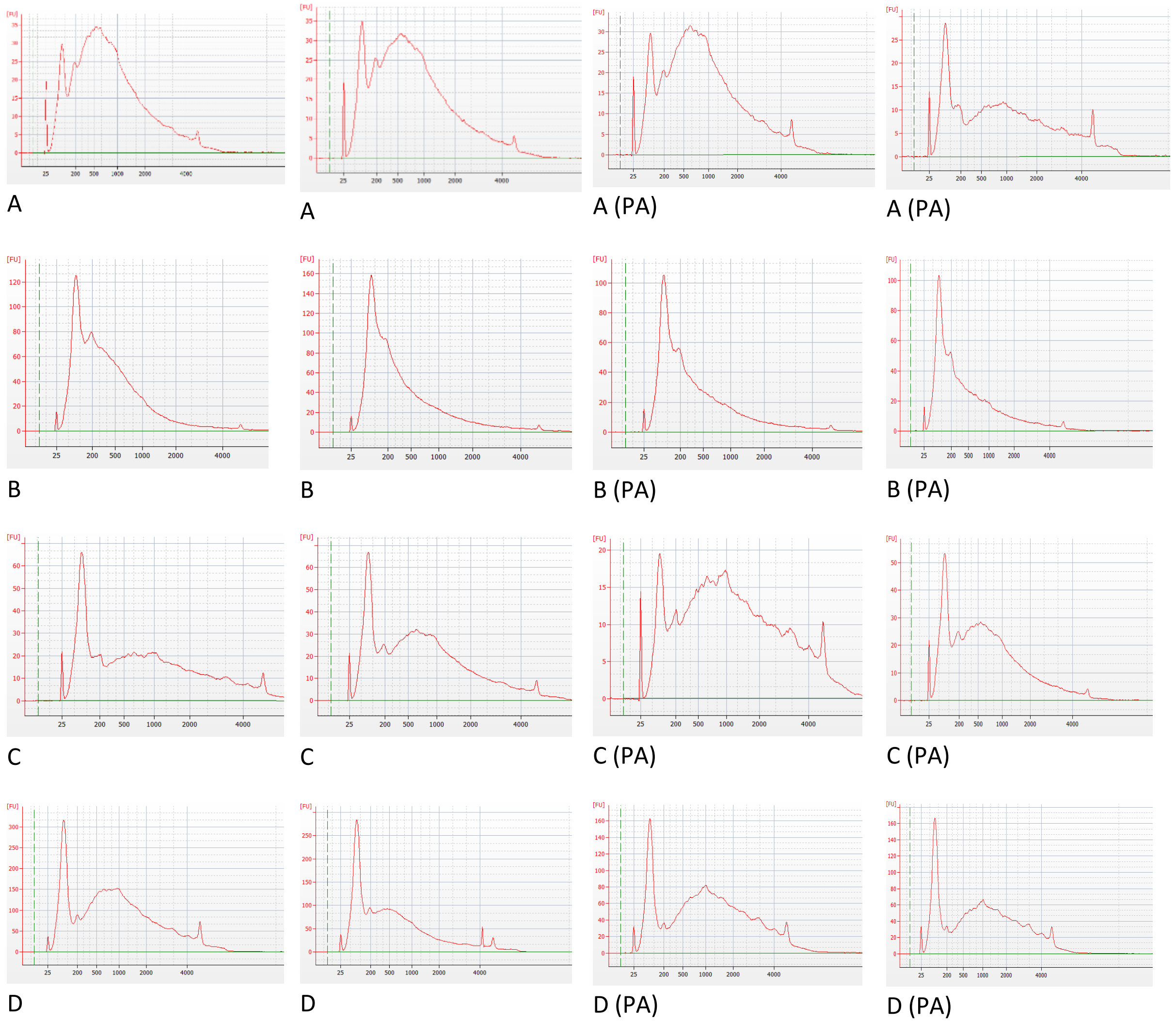

**Figure S3.** Read length distributions for each replicate for cDNA (direct) and PCR-cDNA (PCR) kits for 4 *E. coli* strains (A-D). The red dotted line shows the position of the median.

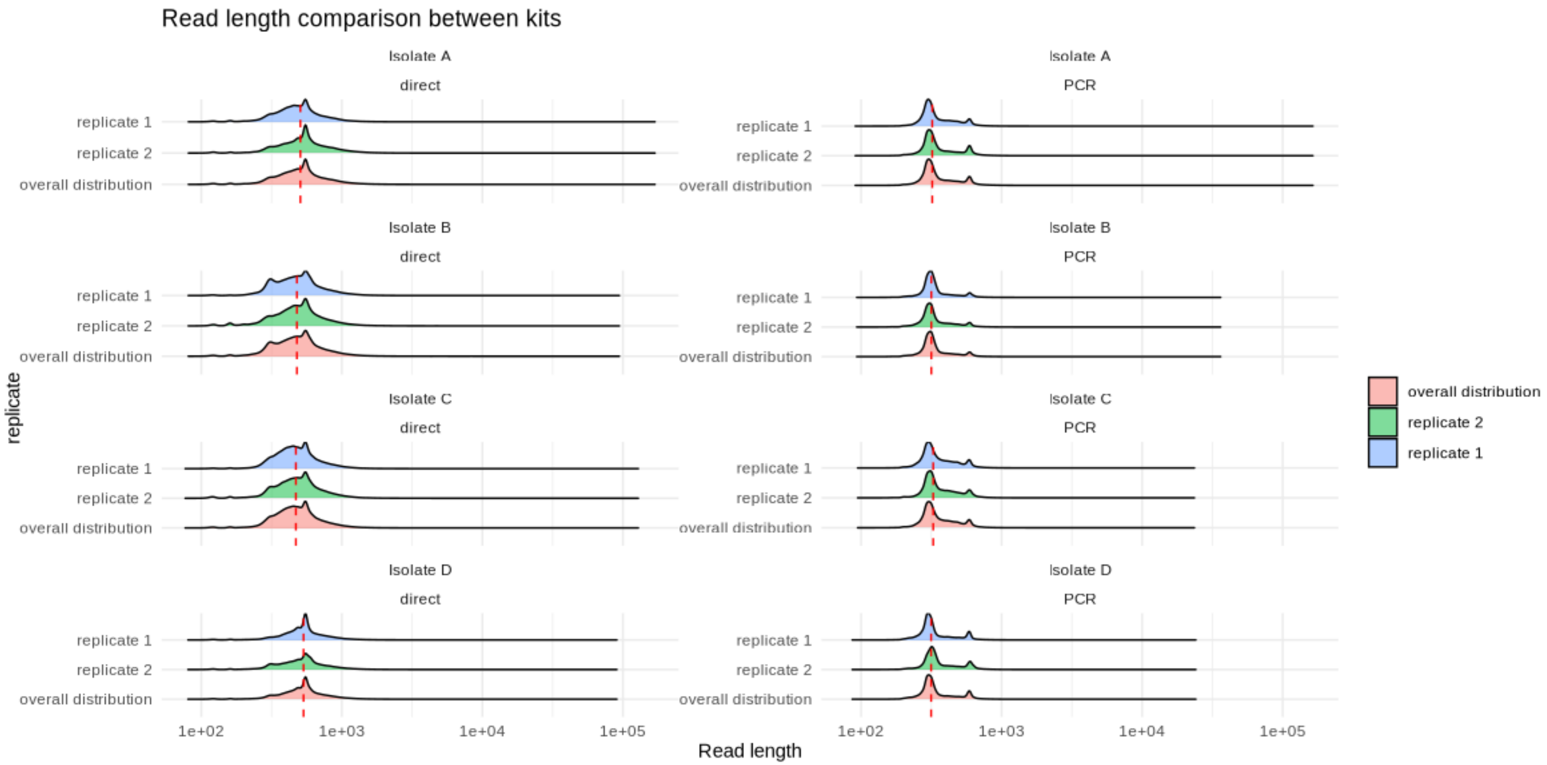

**Figure S4** Read quality distributions for each replicate for cDNA (direct) and PCR-cDNA (PCR) kits for 4 *E. coli* strains (A-D). The red dotted line shows the position of the median.

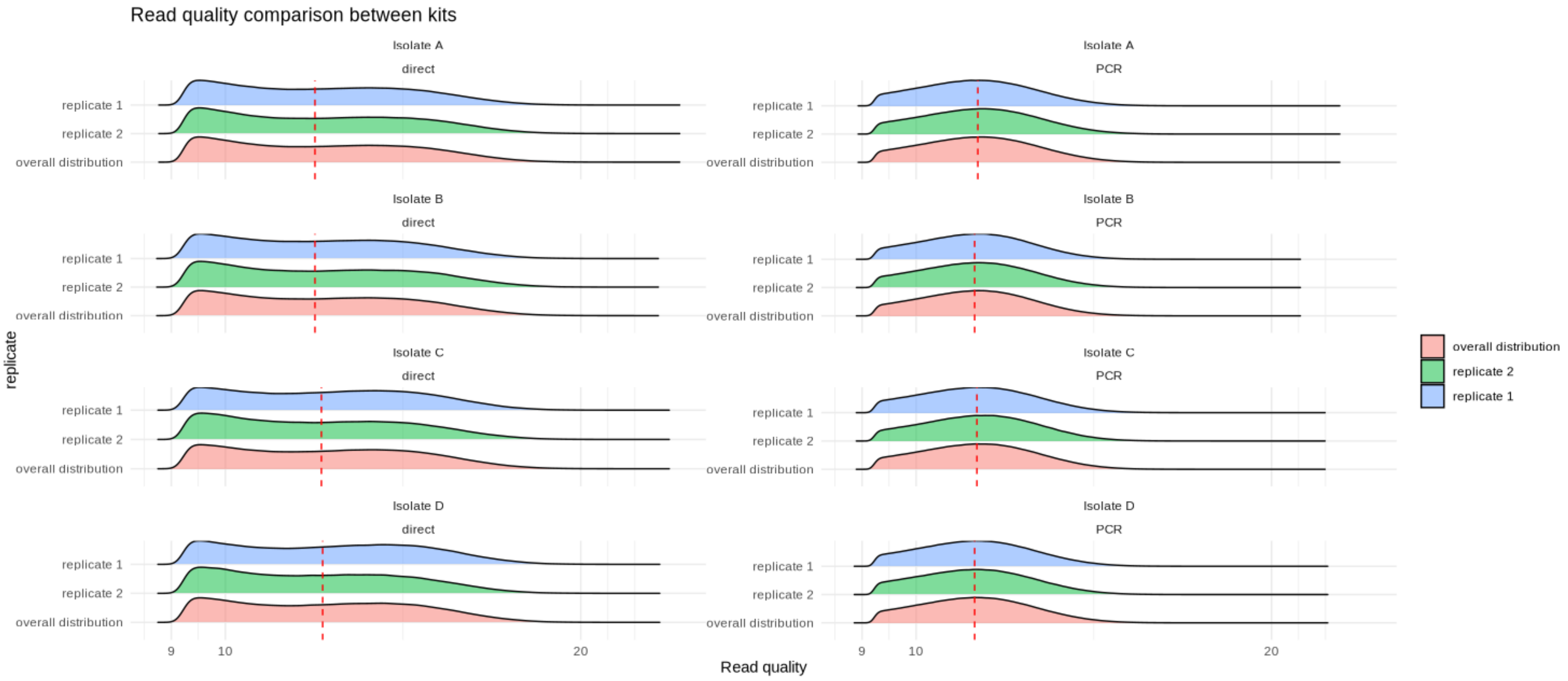

**Figure S5** - Comparison of percentage of reads mapped to reference transcripts between isolates and kits.

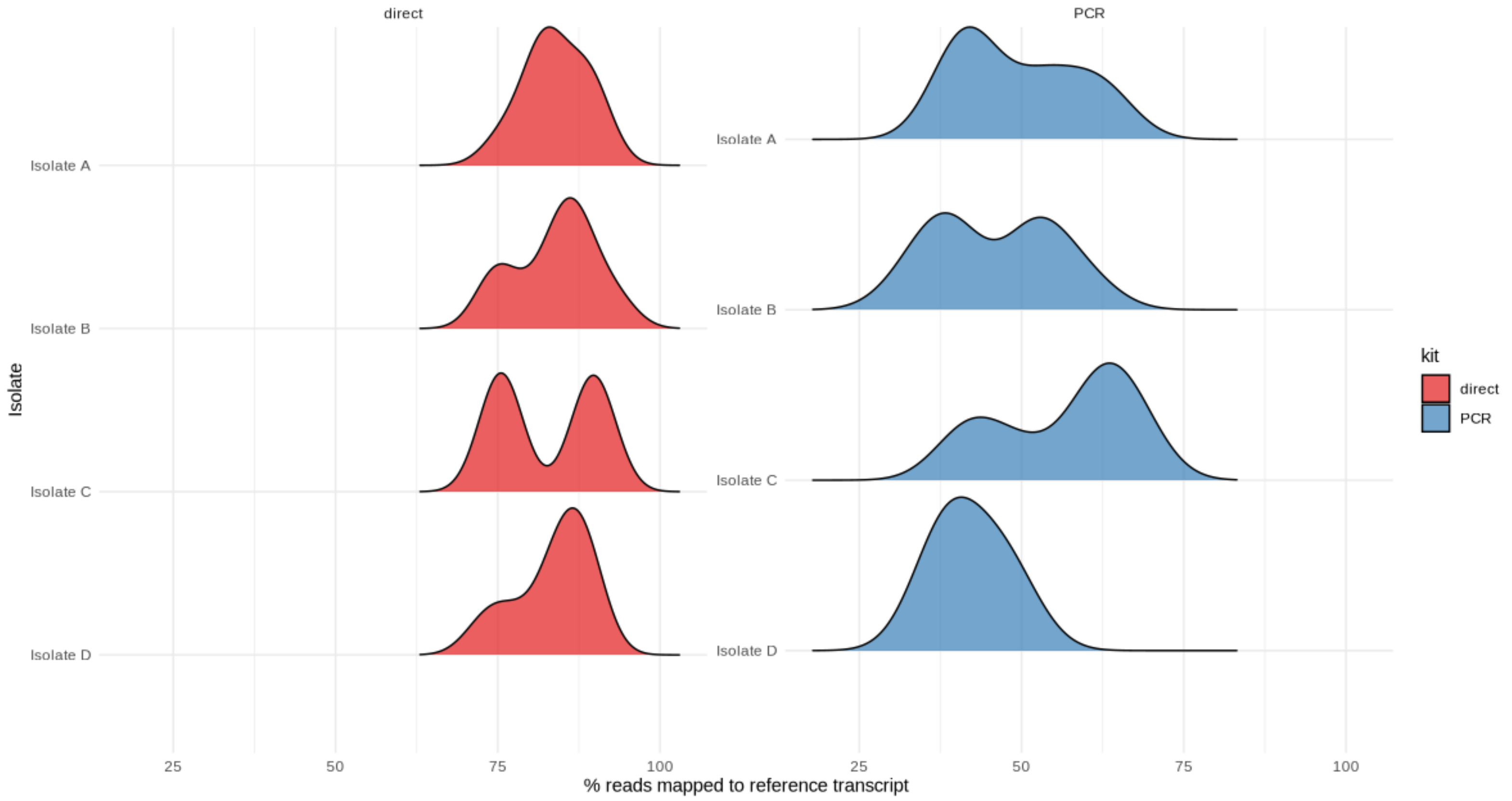

**Figure S6** - Comparison of quality scores (left panel) and read lengths (right panel) for mapped versus unmapped reads between kits (i.e. direct versus PCR).

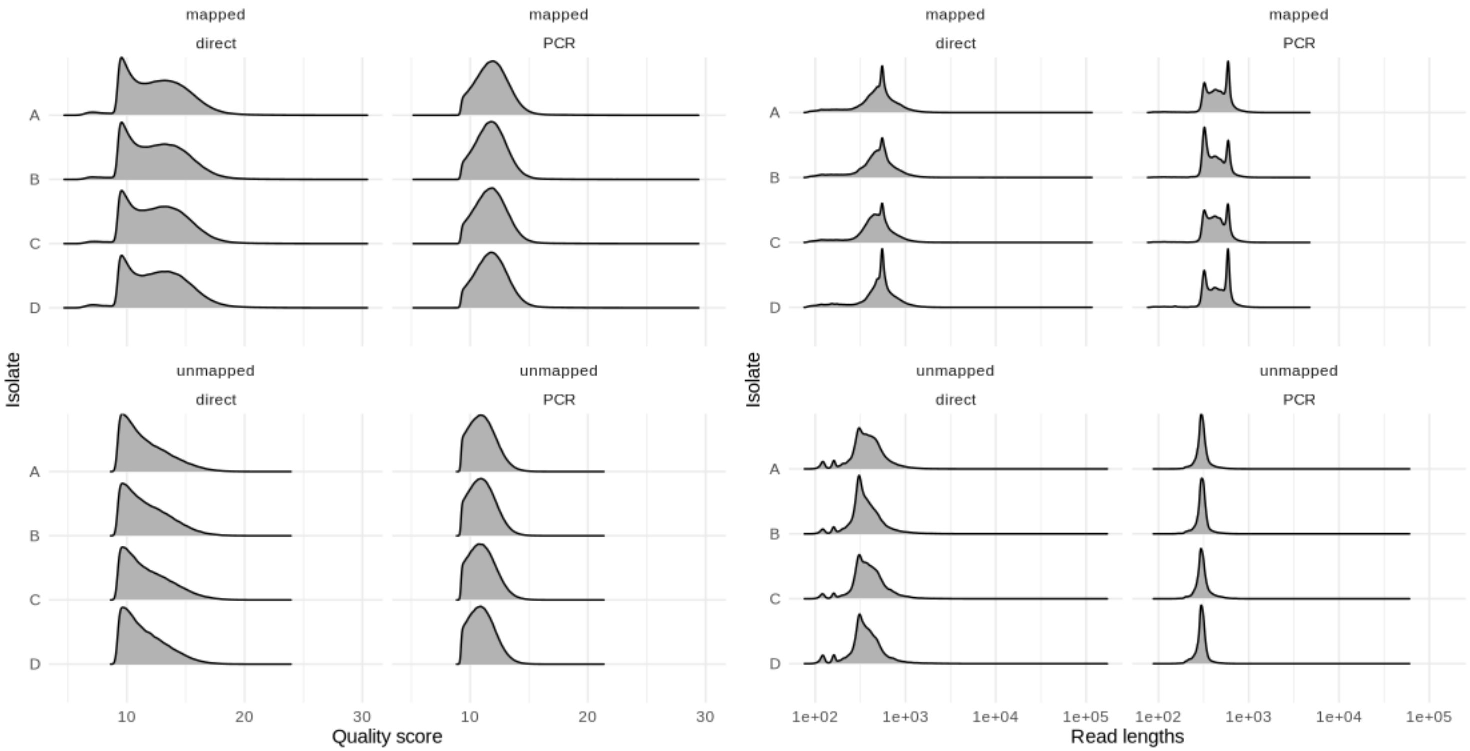

**Figure S7**. Correlogram showing correlations between read counts at a gene level for combination of isolates A_/B_/C_/D_, replicates r1/r2, kits PCR/direct and flowcells 1/2/3/4.

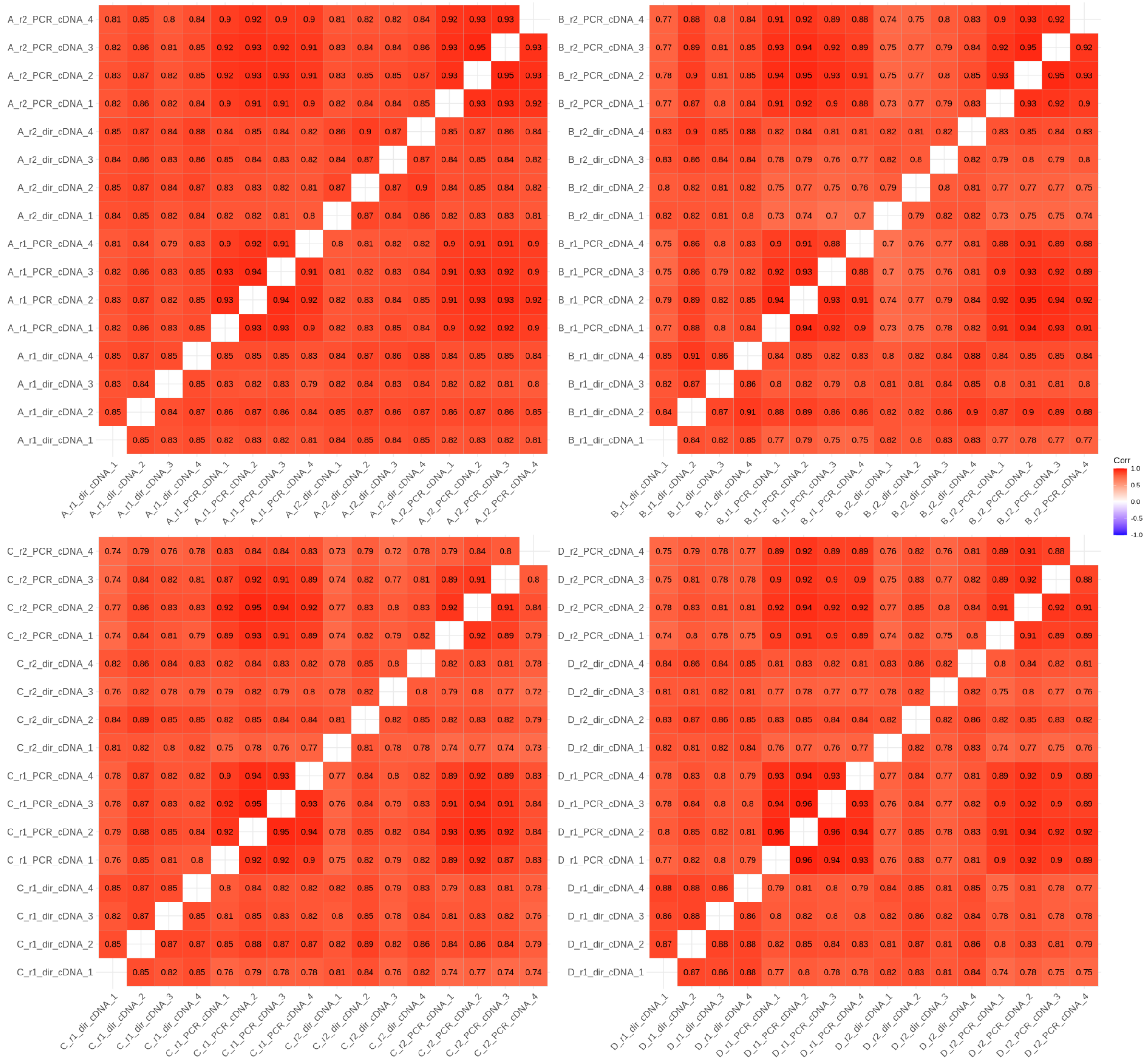

1. Pitt ME, Nguyen SH, Duarte TPS, Teng H, Blaskovich MAT, Cooper MA, et al. Evaluating the genome and resistome of extensively drug-resistant Klebsiella pneumoniae using native DNA and RNA Nanopore sequencing. Gigascience. 2020;9(2).
